# The impact of tumor receptor heterogeneity on the response to anti-angiogenic cancer treatment

**DOI:** 10.1101/218073

**Authors:** Ding Li, Stacey D. Finley

**Author notes:** Corresponding author: Stacey D. Finley, 1042 Downey Way, DRB 140, Los Angeles, CA 90089.

## Abstract

Multiple promoters and inhibitors mediate angiogenesis, the formation of new blood vessels, and these factors represent potential targets for impeding vessel growth in tumors. Vascular endothelial growth factor (VEGF) is a potent angiogenic factor targeted in anti-angiogenic cancer therapies. In addition, thrombospondin-1 (TSP1) is a major endogenous inhibitor of angiogenesis, and TSP1 mimetics are being developed as an alternative type of anti-angiogenic agent. The combination of bevacizumab, an anti-VEGF agent, and ABT-510, a TSP1 mimetic, has been tested in clinical trials to treat advanced solid tumors. However, the patients’ responses are highly variable and show disappointing outcomes. To obtain mechanistic insight into the effects of this combination anti-angiogenic therapy, we have constructed a novel whole-body systems biology model including the VEGF and TSP1 reaction networks. Using this molecular-detailed model, we investigated how the combination anti-angiogenic therapy changes the amounts of pro-angiogenic and anti-angiogenic complexes in cancer patients. We particularly focus on answering the question of how the effect of the combination therapy is influenced by tumor receptor expression, one aspect of patient-to-patient variability. Overall, this model complements the clinical administration of combination anti-angiogenic therapy, highlights the role of tumor receptor variability in the heterogeneous responses to anti-angiogenic therapy, and identifies the tumor receptor profiles that correlate with a high likelihood of a positive response to the combination therapy. Our model provides novel understanding of the VEGF-TSP1 balance in cancer patients at the systems-level and could be further used to optimize combination anti-angiogenic therapy.

## Introduction

Angiogenesis is a hallmark of cancer that facilitates tumor progression in many aspects (1). Tumor growth relies on the formation of new blood vessels to enable waste exchange and to deliver oxygen and nutrients to the tumor. In addition, angiogenesis increases the likelihood of metastasis by enabling tumor cells to enter the bloodstream and disperse to other sites in the body (2). Considering the outstanding importance of angiogenesis for tumor development, anti-angiogenic therapy was designed to starve the tumor of its nutrient supply and limit its growth (3).

Tumor angiogenesis is controlled by both pro-and anti-angiogenic signaling (4–6). Common anti-angiogenic therapy uses single agents to reduce the pro-angiogenic signals, and the primary anti-angiogenic agent being used in the clinic inhibits signaling mediated by vascular endothelial growth factor-A (VEGF), a potent promoter of angiogenesis. However, this approach is not effective in all cancers. For example, bevacizumab, a monoclonal antibody that binds VEGF, is no longer approved for the treatment of metastatic breast cancer because it was not shown to be effective and safe for patients (7). Sunitinib, a tyrosine kinase inhibitor that targets VEGF receptors and other growth factor receptors, has also shown limited success (8). These limitations of anti-VEGF treatment prompt the need to optimize anti-angiogenic therapy. One alternate approach is to enhance the signal of anti-angiogenic factors. Thrombospondin-1 (TSP1) is one of the most studied endogenous inhibitors of angiogenesis and has been shown to inhibit vascular growth and tumorigenesis in preclinical trials (9–11). Inspired by the effect of TSP1, TSP1 mimetics were developed for tumor treatment (12). One such drug, named ABT-510, reached Phase II clinical trials. However, ABT-510 failed to show clear evidence of efficacy and is no longer tested as a single-agent drug in clinical development (13,14).

The disappointing outcomes of clinical studies of anti-angiogenic drugs as single agents prompt the development of combination anti-angiogenic therapy. Administering a combination of anti-angiogenic agents that simultaneously target multiple angiogenic signals is expected to achieve efficient and durable suppression of angiogenesis by strongly shifting the relative balance of inducers and inhibitors of angiogenesis to oppose the “angiogenic switch” (15,16). The combination of agents targeting different pathways might also prevent tumors from leveraging complementary pathways to escape anti-angiogenic treatment. With this in mind, the ABT-510 TSP1 mimetic was clinically tested in combination with bevacizumab in patients with advanced solid tumors. However, patients displayed a heterogeneous response to this combination therapy (17): one patient had a partial response and only 32% of the patients had prolonged stable disease (≥ 6 months). Unfortunately, the mechanisms driving these disappointing results were not elucidated in the trial. Three fundamental questions remain: do the levels of TSP1 and VEGF balance one another in tumor tissue, how does combination therapy influence this VEGF-TSP1 balance, and does inter-patient heterogeneity lead to significantly different responses. Answering these questions contributes to our understanding of the action of combination therapy. In addition, considering the angiogenic balance at the levels of tissue, organs and the whole body can help us optimize anti-angiogenic therapy. In this study, we address these questions using a computational systems biology model.

Mathematical modeling serves as a useful tool to study the response to anti-angiogenic treatment, complementing experimental and clinical studies. Various modeling approaches have been applied to study anti-angiogenesis therapies, including differential equation based models, boolean network models, multi-compartment models, hybrid cellular automaton models, multiscale agent-based models, image-based models and bioinformatics-based modeling (18). In particular, the effects of anti-VEGF agents on VEGF and its receptors have been intensively investigated with computational modeling. Targeting VEGF binding, VEGF secretion and VEGFR signaling as anti-angiogenic strategy have been illustrated in different studies (18). Recently, mathematical modeling was used to understand the impact of the cross-talk between tumor cells and endothelial cells (19) and the vascular phenotypes (20) on the effects of anti-angiogenic treatment, which generates new insights of anti-angiogenic therapy using mathematical model. In this study, one of our goals is to understand the anti-angiogenic therapy targeting VEGF and TSP1 simultaneously, which has not been the focus of previous models. To achieve this goal, we build a novel ordinary differential equation (ODE) based multi-compartment model.

The model presented here significantly builds upon our previous modeling efforts, and is mainly based on two published models. One model is the whole-body model of the VEGF-receptor system, which was previously used to illustrate the counterintuitive increasing of VEGF after anti-VEGF treatment (21). Second, we build on our recent model of TSP1 and VEGF interactions in tumor tissue, which predicts the effects of various strategies mimicking TSP1’s anti-angiogenic properties (22). This model of the VEGF-TSP1 balance in tumor tissue, however, omits the trafficking of soluble species and only considers drugs delivered directly into the tumor. Here, we significantly expand these two previous works (22,23) to generate a novel whole-body model of the VEGF-TSP1 interaction network. The expanded model has three compartments to incorporate the drug pharmacokinetics (PK) and species distribution in the human body. The model also incorporates pharmacodynamics (PD). Since the binding of angiogenic factors to their receptors triggers a cascade of intracellular reactions, including phosphorylation of the receptors, we use the number of ligand-receptor complexes as an approximation of the receptor activation level to capture the status of pro- and anti-angiogenic signaling in tissue. Altogether, our new model enables a complete PK/PD study of the clinically tested combination anti-angiogenic therapy targeting both VEGF and TSP1.

We apply the model to understand how the angiogenic balance of VEGF and TSP1 is modulated by bevacizumab and ABT-510 combination therapy. Then we use the model to investigate the impact of inter-patient heterogeneity, specifically the tumor receptor heterogeneity, on the response to combination anti-angiogenic therapy. Compared to other inter-patient variability, tumor receptor heterogeneity is one of the most well supported in published literature. It has been observed experimentally that tissue samples from patients with different types of cancer and different samples from patients with the same type of cancer have different receptor expressions (24–27). Additionally, the VEGFR2 heterogeneity was shown to affect the response to an anti-angiogenic cyclophosphamide treatment in an *in vitro* experimental setting (28). The patient-to-patient VEGFR1 and neuropilin variability has been associated with the response to bevacizumab treatment as intra-tumoral biomarkers (29,30). Thus, understanding the effects of tumor receptor variability is clinically relevant (28–31). Our model predicts that the metric of angiogenic receptor expression can serve as a predictive tissue biomarker to distinguish the patients in which the combination anti-angiogenic therapy will elicit a strong therapeutic response. Overall, we establish a new computational framework to predict the effects of anti-angiogenic therapies and understand clinical observations.

## Methods

### The compartmental whole-body model

This mechanistic model characterizes the extracellular distribution of angiogenic species in the human body. We follow the compartmental model structure used in previous works (23,32). In this approach, a tissue is assumed to be a collection of capillaries, surrounded by parenchymal cells. The interstitial space lies between the parenchymal cells and the capillaries, which is comprised of the extracellular matrix (ECM), parenchymal basement membranes (PBM) and endothelial basement membranes (EBM). The soluble species are assumed to diffuse within the available interstitial space very fast compared to the timescale of the biochemical reactions (33), thus all of the structures are modeled in a spatially-averaged manner as a simplification. A human cancer patient is represented by a three-compartment model: normal tissue (“normal,” represented by skeletal muscle), the vasculature (“blood”), and diseased tissue (“tumor”) (***Fig. 1A***). The soluble species are introduced to the system by being secreted by cells and are removed from the system through degradation and clearance from the blood. Receptors are uniformly distributed on the cell surfaces and can be internalized by the cell and recycled back to the surface. The total number for each type of receptor is assumed to be conserved at every simulated time point. Transport of soluble species between the compartments is mediated by transcapillary permeability and lymphatic flow.

**Figure 1.**
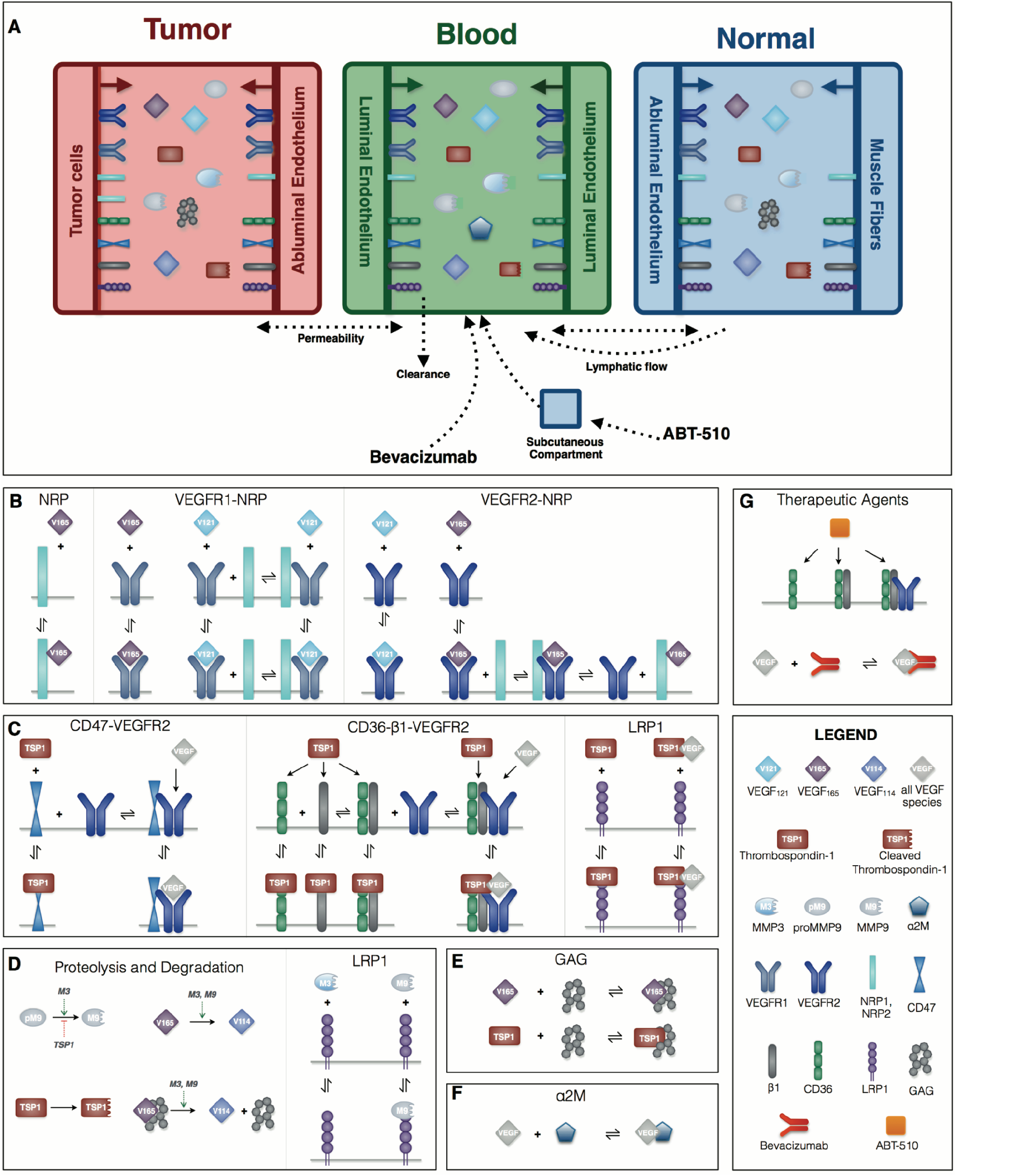
Compartmental model of VEGF-TSP1 system in human cancer patients and TSP1 and VEGF interactions. (A) The model includes three compartments: normal tissue, blood, and tumor tissue. The endothelial cells and parenchymal cells secrete soluble species (VEGF, TSP1, MMP3, and proMMP9) into the compartments. Receptors are localized on the luminal and abluminal surfaces of endothelial and parenchymal cells. Free and ligand-bound receptors can be internalized. Transport between compartments occurs by trans-endothelial permeability and lymphatic flow. Soluble species are degraded in the tissue or cleared from the blood. Specifically, the compartment model includes: (B) The molecular interactions of two active VEGF isoforms (VEGF_121_ and VEGF_165_), receptors (R1 and R2), and co-receptors (N1 and N2); (C) The interactions between TSP1, its receptors (CD36, CD47, β1, and LRP1), VEGF and VEGFR2; (D) The activation of proMMP9 via cleavage by MMP3; cleavage of TSP1; and proteolysis of VEGF_165_ (free or bound to glycosaminoglycan, GAG, chains) to form VEGF_114_ by active MMPs; (E) The sequestration of VEGF_165_ and TSP1 by GAG chains in the extracellular matrix and the cellular basement membranes; (F) The sequestration of VEGF by α2M in blood; (G) The binding between ABT-510 and receptors and sequestration of VEGF by bevacizumab.

Model construction is based on certain assumptions. Firstly, we formulate the compartmental model assuming that the tumor volume is constant. Admittedly, there is a change in the number of healthy or diseased cells in human patients undergoing anti-angiogenic therapy, as a primary goal of treatment is to reduce tumor volume. However, as we have shown in our previous work (34), that since the tumor is nearly 2,000 times smaller than the normal compartment, the tumor volume must change by at least two orders of magnitude (to ∼3300 cm^3^) for it to significantly influence the distributions of the soluble factors, which is a central focus of our PK/PD compartment model. Since this size of tumor is not physiologically realistic, we assume constant tumor volume and instead focus on the distribution of the soluble factors and the formation of pro- and anti-angiogenic complexes in each of the compartments. Accordingly, we assume that the total surface area of the microvessels is constant, as the tissue vascularity is characterized as the ratio of the microvascular surface area to the tumor volume. Finally, we assume that the vascular permeability between compartments is fixed. We implement this simplification because there is a scarcity of quantitative data available to formulate a mathematical equation to capture the relationship between the anti-angiogenic therapy and vascular permeability. Rather than impose further uncertainty in the model, we maintain a constant value for the permeability.

### Rule-based model of VEGF-TSP1 reaction network

The characterization of the species’ dynamics is based on the principles of mass action kinetics and biological transport. BioNetGen, a rule-based modeling approach, is used to construct the model (35). The biological reaction rules are defined in BioNetGen, which automatically generates the set of ordinary differential equations (ODEs) that describe how the species’ concentrations evolve over time. A detailed description of the derivation of the ODE model through BioNetGen is documented in ***Supplementary File S1***, which provides an explanation of the propagation of the rules and reactions in the model generation. Here, we briefly summarize the defined rules that govern the molecular interactions and corresponding reactions (***Fig. 1B-G***). Following our previous works, the model includes two active VEGF isoforms (VEGF_165_ and VEGF_121_). The inactive form, VEGF_114_, is the product of proteolytic cleavage of VEGF_165_. Two predominant VEGF receptors, VEGFR1 and VEGFR2 (R1 and R2), and neuropilin co-receptors, NRP1 and NRP2 (N1 and N2), are considered (***Fig. 1b***). TSP1 binds to its receptors, CD36, CD47, low density lipoprotein receptor-related protein 1 (LRP1) and α_x_β_1_ intergrins (β1, a generic form representing several species) (***Fig. 1C***). We also include matrix metalloproteinase species (MMP3, MMP9 and proMMP9), which promote VEGF cleavage. TSP1 impedes the activation of MMP9 (***Fig. 1D***) as a means of inhibiting pro-angiogenic signaling. Glycosaminoglycan (GAG) chains reside in the interstitial space, representing the extracellular matrix, as well as in the cellular basement membranes. GAG chains are able to bind and sequester TSP1 and VEGF_165_ (***Fig. 1E***). The α-2-macroglobulin (α2M) species, a protease inhibitor, is confined to the blood compartment, where it binds to VEGF (***Fig. 1F***).

### Model parameterization

There are 157 parameters presented in our model, including geometric parameters, kinetic parameters, receptor numbers, secretion and degradation rates, transport rates, and parameters for the drug properties. The model parameter values are reported in ***Supplementary File S1***, with literature references and described below.

#### Geometric Parameters (27 Parameters)

The geometric parameters characterize the fundamental structure of the model, defining the volume of compartments, the interstitial space volume, and tissue surface areas of endothelial and parenchymal cells. These parameters are based on experimental measurements taken directly from *in vivo* mouse tumor models (32). We assume that the geometric characteristics of xenograft tumors in mice recapitulate human tumors, rather than relying on data from *in vitro* cell culture. The set of geometric parameters has been used in multiple previous studies (21,23,34), and we adopt the parameters without changing their values.

#### Kinetic Parameters (47 Parameters)

The kinetic parameters specify the association and dissociation rates for the binding of molecular species. For the VEGF axis, the kinetic parameters are based on experimental measurements for the biochemical interactions of VEGF and its receptors, which have been implemented in our previous models (23,36). Likewise, the kinetic parameters for TSP1 axis are based on experimental measurements that estimate the rates of interactions between TSP1 and its receptors and other binding partners, which were systematically reported in our published model of VEGF and TSP1 in tumor tissue (22).

#### Receptor Numbers (32 Parameters)

The receptor numbers for the VEGF axis (VEGFR1, VEGFR2 and neuropilin-1 and −2) are taken from quantitative measurements of receptor expression on cells from mouse xenograft studies. The cell surface expression of these receptors was measured via flow cytometry (24,25). For the TSP1 receptor numbers, we referred to the qualitative measurements reported in the Human Protein Atlas (27), assuming “high”, “medium”, and “low” expression levels correspond to 10,000, 5,000, and 2,500 receptors/cell, respectively.

#### Secretion and Clearance Rates (34 Parameters)

The clearance and degradation rates are based on the reported protein half-life values. The secretion rates of VEGF were fit based on modeling *in vivo* population PK data in our previous study (23). The secretion rates of TSP1, MMP3 and proMMP9 (10 parameters) are fitted in this study to match the experimental measurements shown in ***Table 1***.

**Table 1.** Comparison of predicted and experimental steady state concentrations of VEGF, TSP1 and MMPs.

| **Species** | **Range of Experimental Measurements\*** | **Predicted Concentration** | **Source and References** |
| --- | --- | --- | --- |
| **Tumor Tissue** |  |  |  |
| VEGF | 8.0-389 pM | 148.6 pM | Multiple cancer types (34) |
| TSP1<sup>#</sup> | 1.0-6.2 nM (2.0) | 2.0 nM | Breast cancer patients (68) |
| MMP3 | 1.8-65.1 nM (5.1) | 4.9 nM | Oral squamous cell carcinoma (69) |
| MMP9 total | 1.0-287.8 nM (9.0) | 9.3 nM | Oral squamous cell carcinoma (69) |
| MMP9 active | 0-22.4 nM (0.8) | 0.2 nM | Oral squamous cell carcinoma (69) |
| **Plasma** |  |  |  |
| VEGF | 0.4-3.0 pM | 1.6 pM | Multiple cancer types (70) |
| TSP1<sup>#</sup> | 0.8-2.1 nM (1.2) | 1.2 nM | Breast cancer patients (68) |
| MMP3 | 1.9-2.0 nM | 1.9 nM | Breast cancer patients (71) |
| MMP9 total | 0.6-0.7 nM | 0.7 nM | Breast cancer patients (71) |
| **Normal Tissue** |  |  |  |
| VEGF | 0.3-3.0 pM | 1.0 pM | Healthy subjects (34) |
| TSP1<sup>#</sup> | 0.2 nM | 0.9 nM | Healthy subjects (72) |
| MMP3 | 0.9-66.4 nM (4.1) | 4.2 nM | Oral squamous cell carcinoma (69) |
| MMP9 total | 0.8-27.7 nM (5.0) | 3.2 nM | Oral squamous cell carcinoma (69) |
| MMP9 active | 0-4.1 nM (0.02) | 0.05 nM | Oral squamous cell carcinoma (69) |

\* Median value is shown in parentheses, if provided in literature. If the experimental data reflects the total concentration in tissue, we assume 50% of the total protein amount is localized in extracellular space.

#### Transport Rates (6 Parameters)

We assume passive transport for all soluble species in our model. That is, the transport of angiogenic species between compartments is only mediated by lymphatic flow and vascular permeability. The rate of lymphatic flow is based on the reported experimental measurements (37). The vascular permeability of VEGF is determined in our previous work (32), by considering the Stokes-Einstein radius for the VEGF protein. Since other angiogenic factors are similar in size to VEGF, we assume that all of the newly introduced angiogenic species have the same vascular permeability as VEGF. This assumption can be relaxed in future work.

#### Properties of the Anti-angiogenic Drugs (11 Parameters)

The degradation and clearance rates of bevacizumab and ABT-510 are converted from their reported half-life values measured in clinical trial. The vascular permeability of bevacizumab and its binding rates are the same as those used in our previous modeling work (34), which capture the clinically-measured pharmacokinetic data (38) (***Supplementary Fig. S1A***). The vascular permeability of ABT-510 was set to be the same as VEGF. Since ABT-510 is a TSP1-derived peptidomimetic that specifically binds to the CD36 receptor, we assume it has the same affinity to CD36 as TSP1. The bioavailability of ABT-510 is tuned to be 30% in subcutaneous injection (39), in order to match pharmacokinetic data of TSP1 (40) (***Supplementary Fig. S1B***).

In summary, parameters are either taken from our previous modeling studies or estimated based on experimental measurements. Only 11 parameters in total are fitted in this study (the secretion rates of TSP1, MMP3 and proMMP9 in the compartments and the ABT-510 bioavailability). Given the presence of the uncertainty of the parameters, we performed a global sensitivity analysis to understand the robustness of the baseline model predictions (see below).

#### Sensitivity Analysis

We perform the extended Fourier amplitude sensitivity test (eFAST), a global variance-based sensitivity analysis, to understand how the uncertainty of parameters (“model inputs”) affect the baseline model predictions (“model outputs”) (41). We analyzed the effects of three groups of parameters (receptor numbers, kinetic parameters, and vascular permeability) on nine different model outputs (the concentrations of TSP1, VEGF, proMMP9, MMP9, and MMP3; the TSP1-VEGF, proMMP9-MMP3, and MMP3-TSP1 complexes; and the angiogenic ratio) in each of the three compartments. In each case, the parameter values were allowed to vary 10-fold above and below the baseline values (a total range of two orders of magnitude) to account for uncertainty in the model parameters. In the eFAST method, the inputs are varied together, at different frequencies. The Fourier transform of the outputs is calculated to identify the influence of each parameter, based on the amplitude of each input’s frequency. Two different sensitivity indices are generated in the eFAST analysis: the first-order FAST indices, *S_i_*, and the total FAST indices, *S_Ti_*. The first-order indices (*S_i_*) measure the local sensitivity of individual inputs, while the total indices (*S_Ti_*) represent the global sensitivity by accounting for second- and higher-order interactions between multiple inputs. The eFAST method is implemented using MATLAB code developed by Kirschner and colleagues (41). We have performed this analysis to characterize the robustness of our previous models (22,23).

#### Simulating Receptor Variability

To investigate the impact of tumor receptor heterogeneity, we perform a Monte Carlo analysis and vary the receptor expression parameter values in the tumor compartment. In total, the densities of 16 receptors are varied in the simulations: four VEGF receptors on tumor cells (R1_Tum, R2_Tum, N1_Tum, N2_Tum), four TSP1 receptors on tumor cells (CD36_Tum, CD47_Tum, LRP1_Tum, β 1_Tum), four VEGF receptors on tumor endothelial cells (R1_disEC, R2_disEC, N1_disEC, N2_disEC), and four TSP1 receptors on tumor endothelial cells (CD36_disEC, CD47_disEC, LRP1_disEC, β1_disEC). The densities of these receptors were randomly chosen from a uniform distribution within a range of 10-fold above and below the baseline value. We generated 1,000 different combinations of receptor density profiles, representing 1,000 unique cancer patients. We ran the model for each of the receptor profiles with anti-angiogenic treatment to examine how the response to treatment varies across the 1,000 parameter sets.

### Simulation of therapy

#### Administration of Combination Therapy

The combination therapy of bevacizumab and ABT-510 is simulated by mimicking the administration strategy used in clinical trials (17). We first allowed the model to reach steady state (this occurs within 24 hours) before the start of treatment. We then simulated one cycle of the combination therapy: bevacizumab was administered once at the beginning of the cycle; ABT-510 was administered every 12 hours for 14 days. Bevacizumab was given at a dose of 10 mg/kg through intravenous infusion lasting 90 minutes, while ABT-510 was administered at 100 mg twice daily through subcutaneous injection. The bolus of ABT-510 was given directly to a subcutaneous compartment (42) (assumed to be a reservoir with a volume of 30 cm^3^), and it is subsequently transported into blood. The transportation between the subcutaneous and blood compartments is unidirectional and is assumed to occur at the same as the rate transport between the normal and blood compartments. ABT-510 binds to TSP1 receptor CD36 to induce an anti-angiogenic signal (43) (***Fig. 1G***). Bevacizumab is a VEGF antibody that sequesters VEGF to keep it from binding to its receptors, thereby inhibiting the pro-angiogenic signal (***Fig. 1G***).

#### Characterization of the Response to Treatment

In our study, the response to anti-angiogenic treatment is characterized based on the angiogenic balance in tumor tissue. We define the angiogenic balance as the ratio of the concentrations of the pro-angiogenic complexes to the anti-angiogenic complexes. The ratio indicates the activation level of the pro-angiogenic receptors relative to the activation level of anti-angiogenic receptors. Specifically, the pro-angiogenic complexes include the ligand-bound VEGF receptors that are not interacting with active TSP1 receptors. Here we assume a ligand-bound VEGF receptor coupled with active TSP1 receptor is not a pro-angiogenic complex, since the downstream signaling of the ligand-bound VEGF receptor could be inhibited by TSP1 (44,45). The anti-angiogenic complexes are the active TSP1 receptors, those bound to TSP1 or the TSP1 mimetic. The fold-change of the angiogenic ratio in the tumor compartment (*F*) characterizes the response to anti-angiogenic treatment:

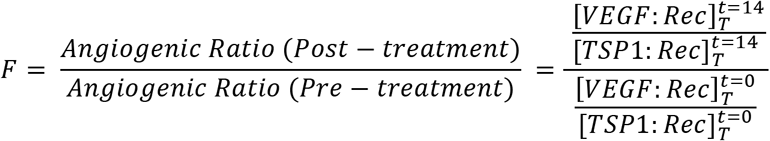

where [*VEGF: Ree*] is the total concentration of pro-angiogenic complexes, [*TSP1-.Rec*] is the total concentration of anti-angiogenic complexes, *T* denotes Tumor compartment, and *t* indicates the simulated time: before treatment (*t* = 0 day) and after one cycle of treatment (*t* = 14 days).

### Quantification of combination effect

We determine the combined effect of the two anti-angiogenic agents, bevacizumab and ABT-510. Commonly used dose–effect-based approaches to quantify drug combination effects rely on the mathematical framework known as Loewe Additivity (46). It calculates the combination of dose *a* for drug *X* and dose *b* for drug *Y* that can produce the same effect as dose *A* of drug *X* alone and dose *B* of drug *Y* alone. The combination effect of drug *X* and drug *Y* can be expressed as:

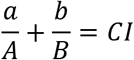

where *CI* is the combination index. Additionally, reference drug values are usually set based on a desired end-point (i.e., the IC50 value). However, here, there is no specified characteristic of the efficacy of the anti-angiogenic drugs (individually or in combination). Therefore, we selected a specific end point (a desired “fold-change of angiogenic ratio”, 0.75). This end point is firstly used to determine the drug doses *a* and *b* in combination. The ratio between doses *a* and *b* is a constant, equal to the ratio used in clinical protocol. Then, we change the doses of bevacizumab and ABT-510 to get the dose-effect curve of single-agent treatments and find doses *A* and *B* corresponding to the desired end point 0.75. A *CI* value less than one indicates synergy between the two drugs, a *CI* equal to one represents simple additivity, and *CI* greater than one indicates antagonism (47).

### Partial least squares regression

Partial least squares regression (PLSR) relates input variables to output variables by maximizing the correlation between the variables. This is accomplished by projecting the output variables onto new dimensions (principal components, PCs), which are linear combinations of the inputs. We use PLSR to investigate the relationship between tumor receptor numbers and the response to anti-angiogenic therapy (the fold-change in the angiogenic ratio in the tumor tissue, termed *F*). The PLSR model is trained and validated with the 1,000 different combinations of tumor receptor densities (inputs) and the corresponding responses to treatment generated by the three-compartment systems biology model (outputs). The receptor values are normalized by dividing by the lower bound of the sampling range. Since the receptor numbers are sampled from a uniform distribution within 10-fold above and below the baseline value, the normalized receptor values range from 1 to 100. The nonlinear iterative partial least squares (NIPALS) algorithm was used to implement the PLSR analysis (48). Leave-one-out cross validation is applied to quantify the predictive capability of the PLSR model. Furthermore, we calculated the “variable importance of projection” (VIP) for each input variable (49), which is the weighted sum of each input’s contribution to the output. As such, the VIP evaluates the overall importance of an input variable in predicting the output. VIP values greater than one indicate variables that are important for predicting the output response.

## Results

### Baseline model is calibrated to experimental data

We report the species’ concentrations (VEGF, TSP1, MMP3 and MMP9) predicted by the baseline model and compare it with experimental measurements in ***Table 1***. Our predictions quantitatively match the experimental data, where all of the predicted concentrations are within the measured ranges. We also compare the predicted pharmacokinetics of bevacizumab with the data collected in clinical trials (***Supplementary Fig. S1*)** and confirm that our model can capture the dynamics of these two agents at different dosages.

We performed a global sensitivity analysis to quantify the robustness of the baseline model predictions. The eFAST method is used to calculate the sensitivity indices *S_i_* and *S_Ti_* (see Methods section). The *S_i_* estimates how an input influences an output individually, while the *S_Ti_* indicates how influential an input is in combination with other parameters. We find that the predicted species’ concentrations in tumor compartment are mainly affected by the densities of receptors on tumor cells and diseased endothelial cells (***Supplementary Fig. S2A***). Additionally, the predicted concentrations for species in the blood and normal compartments are heavily affected by the receptor densities on normal endothelial cells and normal parenchymal cells. The vascular permeability significantly affects the model predictions (***Supplementary Fig. S2B***). Intuitively, the predicted concentrations in the tumor compartment are mainly affected by the permeability between tumor and vascular system, while the model predictions for blood and normal compartment are predominantly affected by permeability between normal tissue and vascular system. The kinetic parameters are shared by reactions in all three compartments, thus they influence the predictions in all three compartments to varying degrees (***Supplementary Fig. S2C***).

Overall, the eFAST results indicate the parameters that should be fitted or for which additional experimental data is needed. Vascular permeability is known to be affected by VEGF and anti-angiogenic drugs. However, to our knowledge, there are no robust, quantitative measurements available that can be used to specify the mathematical relationship between VEGF or anti-angiogenic agents and permeability. Therefore, we keep the permeability constant, based on the size of the molecular species (see Methods). All of the kinetic parameter values, with the exception of the coupling rates between CD36 and VEGFR2, CD36 and β1, and CD47 and VEGFR2, are based on the reported experimental measurements. The three coupling rates for which we do not have experimental values are estimated to have little influence on the model predictions: kc_CD36:R2, kc_CD36:β1 and kc_CD47:R2 are either not significant or have very low *S_i_* and *S_Ti_* values (***Supplementary Fig. S2C***, last three columns). Therefore, the uncertainty of those parameters does not hamper the robustness of the predictions.

Overall, the sensitivity analysis demonstrates that the model is mostly affected by parameters whose values can be specified from experimental data. Below, we present results from applying the calibrated model to predict the effects of anti-angiogenic therapy. We explicitly study the effects of variability in the tumor receptor numbers, as those are shown to influence the predicted concentrations of the angiogenic factors, both in our sensitivity analysis and in experimental and clinical studies (28–31).

### The angiogenic balance is predicted to shift during combination therapy

We apply the compartmental model to predict how the angiogenic signal changes with anti-angiogenic treatment. We simulate one cycle (14 days) of combination therapy, following the protocol used in clinical trial (17). Bevacizumab was given at a dose of 10 mg/kg through intravenous infusion lasting 90 minutes. ABT-510 was administered at 100 mg every 12 hours for 14 days. The concentrations of unbound (“free”) VEGF and TSP1 over time, along with dynamics of the “angiogenic ratio”, are reported to show the change of angiogenic signals during combination therapy (***Fig. 2***). The angiogenic ratio is the number of pro-angiogenic complexes over the number of anti-angiogenic complexes (see Methods). If the angiogenic ratio is greater than one, the compartment is considered to be in a pro-angiogenic state.

**Figure 2.**
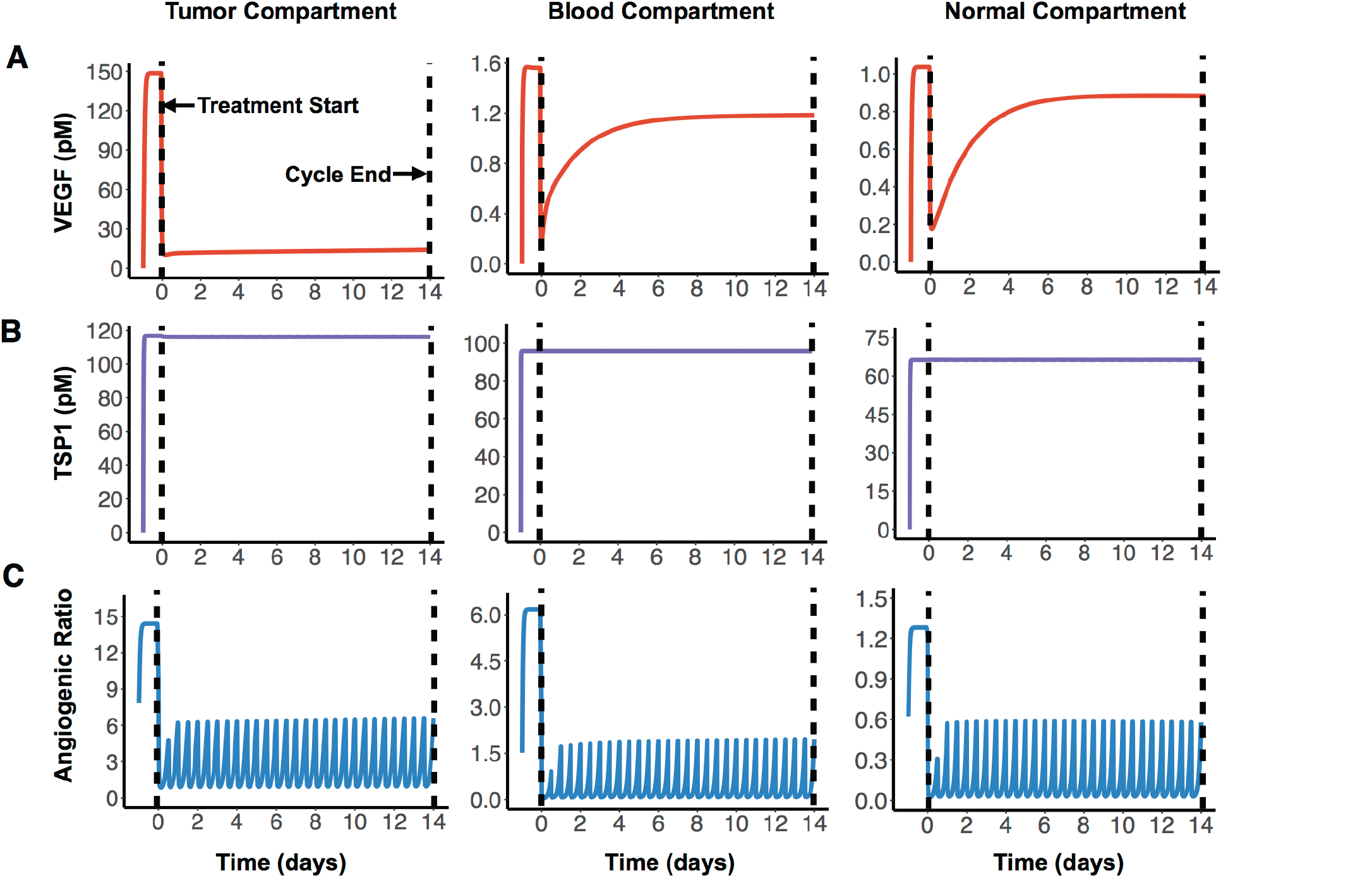
The effects of one cycle of combination anti-angiogenic therapy. The system is allowed to reach steady state before the start of treatment (Day 0). One cycle of treatment lasts 14 days: bevacizumab is given once every two weeks, and ABT-510 is given twice daily. (A) VEGF is shown to decrease after the treatment start in all three compartments. (B) TSP1 is predicted to stay at a stable level. (C) The angiogenic ratio (the ratio of pro-angiogenic complexes to anti-angiogenic complexes) is predicted to oscillate throughout the course of treatment.

After one cycle of the combination therapy, the concentration of free VEGF is predicted to decrease in all three compartments (***Fig. 2A***). The tumor VEGF level changes from 149 pM to 14 pM after one treatment cycle. The levels of free VEGF in plasma and normal tissue decreased from 1.6 pM to 1.2 pM and from 1.0 pM to 0.9 pM, respectively. In contrast, the free active TSP1 level in all three compartments is highly stable throughout the whole treatment cycle (***Fig. 2B***), remaining at 117 pM, 96 pM, and 66 pM in the tumor, blood, and normal compartments, respectively. This indicates that the combination therapy does not affect the free TSP1 level in human body. The experimental measurements from clinical study also show that the level of TSP1 does not significantly change (17), which validates this prediction. However, the combination therapy is predicted to significantly reduce the angiogenic ratio in all compartments. Before treatment, the angiogenic ratio (***Fig. 2C***) is predicted to highly favor angiogenesis in tumor and blood, where the ratios are 14.4 and 6.2, respectively. In comparison, the angiogenic ratio is almost balanced in the rest of the body (1.3 in normal compartment). The values of the angiogenic ratios at the end of the treatment cycle are 6.6, 2.0, and 0.6 in the tumor, blood, and normal compartments. Thus, following treatment, the angiogenic balance is significantly reduced, opposing angiogenesis in normal tissue, but still favoring angiogenesis in tumor tissue and blood. In all compartments, the angiogenic ratio is predicted to oscillate during the treatment. Since the ABT-510 drug has a short half-life (1.2 hours in circulation), it requires a high frequency of administration (twice-daily injections), which causes the angiogenic ratio to oscillate following each injection. Specifically, the angiogenic ratio decreases to a low level for a short time after the administration of ABT-510 and goes back to a medium level before the next dose of ABT-510.

To get detailed insight into how the angiogenic ratio varies in the tumor, we also report the change of all tumor angiogenic complexes in combination therapy and single agent treatment (***Supplementary Fig. S3***). VEGF can bind to four different receptors, including VEGF receptor 1 (R1), VEGF receptor 2 (R2), neuropilin-1 (N1) and neuropilin-2 (N2). Thus, four different types of pro-angiogenic complexes are formed. TSP1 binds to four different receptors, including CD47, CD36, LRP1 and β1, to produce four types of anti-angiogenic complexes. Our predictions show that bevacizumab promotes a decrease in the pro-angiogenic complex involving VEGFR1 (***Supplementary Fig. S3A***), and ABT-510 drives an increase in the anti-angiogenic complex involving CD36 (***Supplementary Fig. S3B***). The absolute levels of other angiogenic complexes only show small changes. In ***Supplementary Fig. S3C***, the curve shows aspects of the curves in ***Supplementary Figs. S3A-B***, which indicates that the result of these two agents in combination involves each of the drug’s individual effects.

### The therapeutic response varies due to tumor receptor heterogeneity

To examine the impact of tumor receptor heterogeneity, we varied the number of VEGF and TSP1 receptors in the tumor tissue, both on tumor cells and diseased endothelial cells. Our sensitivity analysis shows that these receptor numbers significantly influence the tumor angiogenic ratio. Additionally, pre-clinical and clinical investigations indicate that tumor receptors affect anti-angiogenic treatment (28–31). We set the density of each receptor by sampling within two orders of magnitude of the baseline value. This variability in the receptor number influences the dynamics of free VEGF, free TSP1, and the angiogenic ratio, as shown in ***Supplementary Fig. S4***. We use the “fold-change” of free VEGF, free TSP1, and the angiogenic ratio to characterize the response to combination treatment. The fold-change is the post-treatment level compared to the pre-treatment level. When the fold-change is less than one, the quantity has decreased after treatment. We further define a therapeutic response to be when the fold-change of free VEGF or the angiogenic ratio is less than one, indicating a shifting of the angiogenic balance to oppose angiogenesis. The model predicts that combination treatment elicits a therapeutic effect for all of the receptor levels simulated. That is, the fold-change of free VEGF and the angiogenic ratio are predicted to be lower than one in all three compartments (***Fig. 3A***), indicating that free VEGF and the angiogenic ratio decrease after treatment.

**Figure 3.**
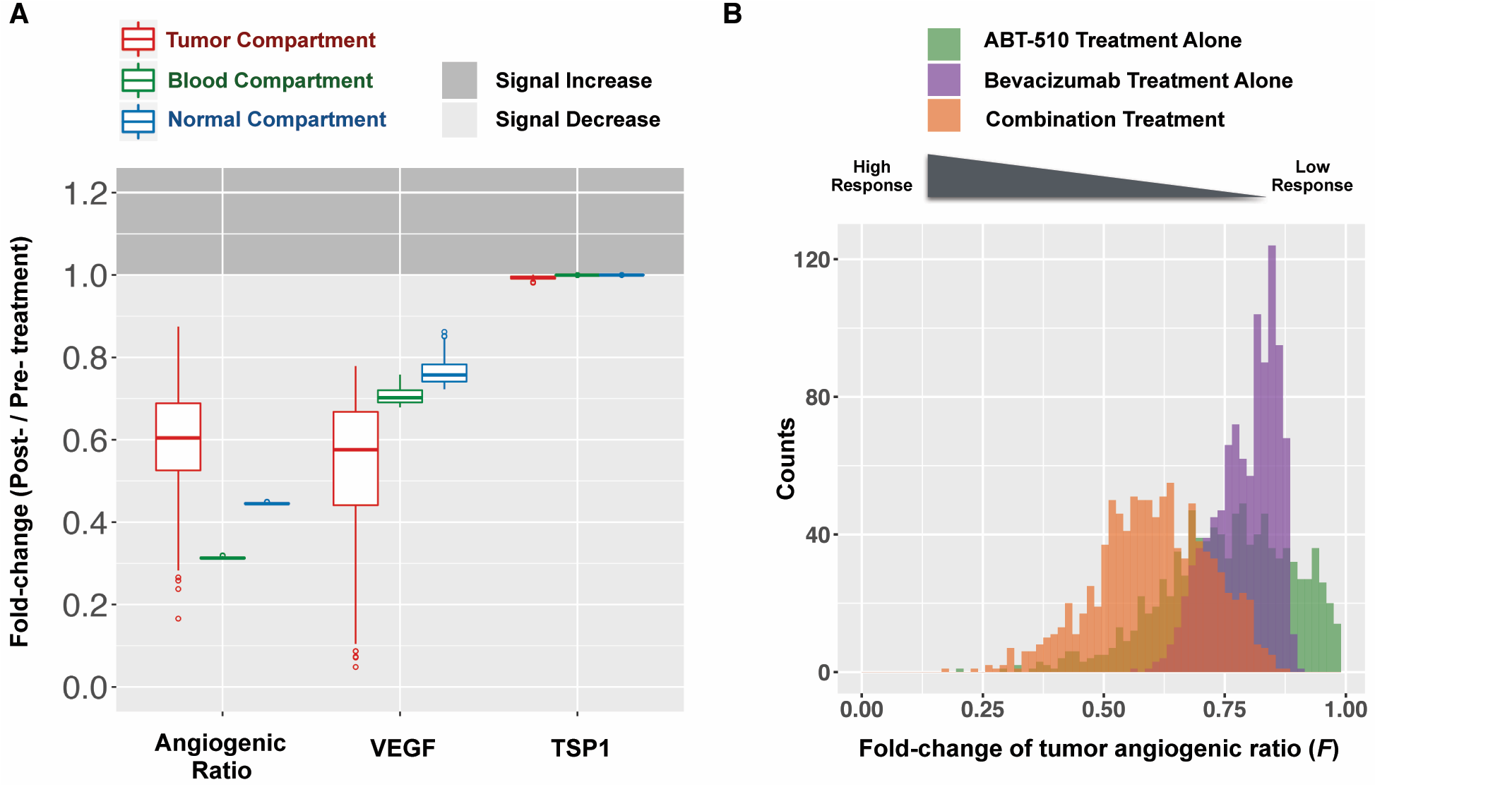
The impact of tumor receptor heterogeneity on the response to anti-angiogenic therapy. We simulated 1,000 tumor receptor profiles and predict: (A) The fold-changes of the angiogenic ratio, VEGF, and TSP1 in tumor (red), blood (green), and normal (blue) compartments. (B) The distributions of the fold-change of the tumor angiogenic ratio (*F*) for single-agent treatments and combination therapy (*green*: ABT-510 alone; *purple*: bevacizumab alone; *orange*: combination treatment).

Model predictions show that the response level of the fold-change in tumor varies depending on the tumor receptor profile. Different combinations of sampled parameter values represent patients with different tumor receptor profiles. The model predicts a wide range of fold-changes for the angiogenic ratio and free VEGF in the tumor compartment (***Fig. 3A***), which indicates that different tumor receptor profiles lead to different responses to the combination therapy. Furthermore, the skewed distribution of the fold-changes of the tumor angiogenic ratio (*F*) (***Fig. 3B, orange***) shows that the majority (83%) of the sampled tumor profiles lead to intermediate responses with fold-change ranging from 0.5 to 1. A smaller percentage of profiles (17%) have a strong response to treatment (*F* < 0.5). Interestingly, there are some outliers for which combination treatment significantly decreases the pro-angiogenic signal, with fold-changes of free VEGF or angiogenic ratio much lower than one. These results imply that particular tumor receptor profiles could have a strong therapeutic response to combination therapy. In single-agent anti-angiogenic tumor therapy (ABT-510 or bevacizumab treatment alone), heterogeneous responses are also observed (***Fig. 3b, green and purple***). Thus, the response to single-agent treatment also varies depending on the tumor receptor profile.

Tumor receptor heterogeneity influences treatment outcomes in other aspects. Firstly, variability in the tumor receptor numbers could affect the targeting of combination anti-angiogenic therapy. ***Fig. 3A*** shows the fold-changes in the angiogenic ratio in the blood and normal compartments are ∼0.3 and ∼0.45, respectively. In comparison, in the tumor, the fold-change varies from ∼0.3-0.9, with the median being 0.6. That is, across all 1,000 simulations, the median effect of the combination therapy is a reduction of the angiogenic ratio by 40% in tumor, while the angiogenic ratio is reduced by 70% in blood and by 55% in normal tissue. Thus, for most cases, the fold-change of the angiogenic ratio in healthy tissue and in blood is lower than in the tumor (***Fig. 3a***). This indicates the combination therapy is not always efficiently targeting tumor angiogenesis. That is, the combination therapy is targeting physiological angiogenesis instead of tumor angiogenesis. This insight is not captured by solely examining the fold-change of VEGF.

Secondly, tumor receptor heterogeneity influences the effect of combination therapy on individual angiogenic pathways. The model predicts that the fold-change of the VEGFR2- and neuropilin-1-containing pro-angiogenic complexes in tumor could be higher than one for some sampled cases (***Supplementary Fig. S5***). This indicates that for some tumor receptor profiles, the pro-angiogenic signaling pathway of VEGFR2 or NRP-1 will be enhanced after treatment instead of inhibited. Similar results are seen when simulating bevacizumab treatment (data not shown). Interestingly, these unintended outcomes following bevacizumab treatment predicted by the model have been observed clinically: a substantial increase in phosphorylated VEGFR2 following bevacizumab treatment (50) and variable changes in phosphorylated VEGFR2 in the clinical trial of the combination anti-angiogenic therapy (17).

### Tumor receptor heterogeneity affects the drug combination effect

We used the model to compare the response to single- and dual-agent therapy and find that the response to combination therapy is related to the response to single-agent treatment (***Fig. 4a***). The color change from red to yellow along the diagonal direction (from upper right to lower left) shows that the response to combination treatment (*Fc*) improves when the response to either single-agent treatment is higher. This indicates that a better response to single-agent treatment may correspond to a better response to combination therapy. We then examined the superiority of the drug combination compared to the single agents. The color of the points in ***Fig. 4b*** indicates the difference between the response to combination therapy and the best response amongst the two single-agent treatments. Although this difference changes depending on the tumor receptor profile, the difference is higher than zero for all sampled cases. This indicates that the response to combination therapy is better than single agent treatment. Thus, when considering the fold-change of tumor angiogenic ratio (*F*), the combination therapy of bevacizumab and ABT-510 is superior to single-agent therapy for all of the sampled tumor receptor profiles.

**Figure 4.**
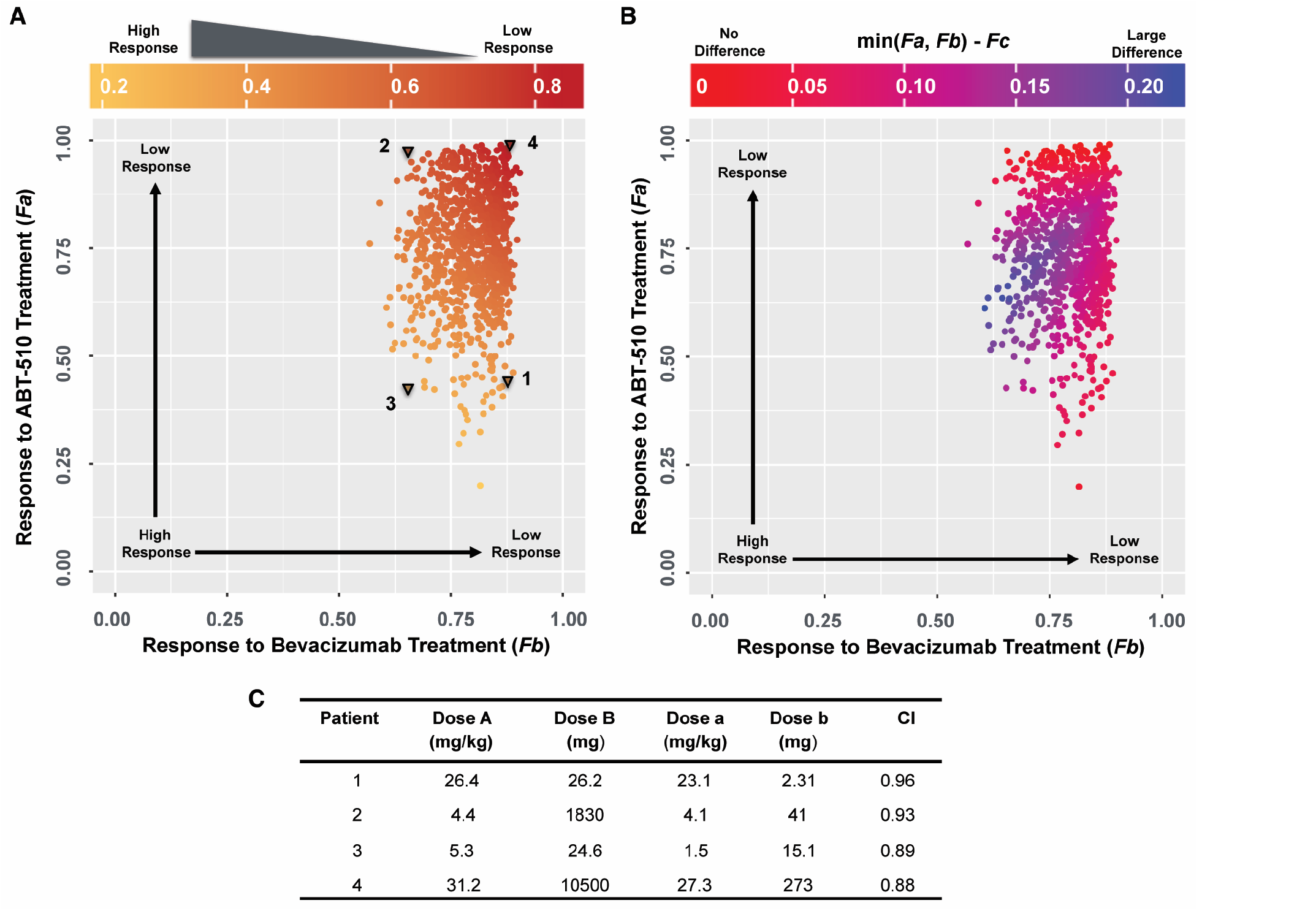
The impact of tumor receptor heterogeneity on the combination effect. (A) The comparison of the response to combination therapy (*Fc*) and responses to single agent treatments (*Fa*: response to ABT-510 alone; *Fb*: response to bevacizumab alone). (B) The comparison of the response to combination therapy (*Fc*) and the best response between the individual treatments (the smaller of *Fa* and *Fb*). (C) The calculated Combination Index (*CI*) for four representative tumor receptor profiles.

We explicitly calculated the combination effect to determine if the action of the two drugs is synergistic or additive. We selected four sets of tumor receptor profiles (***Fig. 4A***, triangles) and calculated the combination index (*CI*) of the Loewe additivity (see Methods). These are representative cases corresponding to patients that respond well to ABT-510 alone (“1”), bevacizumab alone (“2”), or combination therapy (“3”), and one patient that does not respond well to either drug or the combination (“4”). The calculated *CI* values are only slightly different for these four tumor profiles (***Fig. 4C***), where the combination effect is nearly additive for all four patients (*CI* ranges from 0.88 to 0.96). Thus, the bevacizumab and ABT-510 have an additive effect in shifting the tumor angiogenic balance for the four selected patients.

### The therapeutic response depends on tumor cell VEGR1, CD36 and CD47 expression

Given the observed impact of tumor receptor heterogeneity in the predictions from the mechanistic model, we explored if we can use tissue biomarkers (tumor receptor levels) to determine the response to combination anti-angiogenic therapy *a priori*. We used partial least squares regression (PLSR) to characterize the association between tumor receptor levels and the response to combination therapy (the fold-change in the angiogenic ratio in the tumor). Specifically, we established a predictive PLSR model that estimates the response to combination therapy using the tumor angiogenic receptor profile (***Supplementary Fig. S6a***). We used 18 variables as the inputs in PLSR modeling: the 16 VEGF and TSP1 receptor numbers on diseased endothelial cells and tumor cells that were varied and two nonlinear terms comprised of ratios of receptor numbers. The fold-changes estimated by the PLSR model closely match the actual values given by the mechanistic model (the points lie along the 45°-angle line). The high Q^2^Y value of 0.94, indicates that in leave-one-out cross validation, the PLSR model predicts over 94% of the total variation in the fold-change of tumor angiogenic ratio. The R^2^Y value of 0.93 shows the most of the variation in the response is also captured by the PLSR model, while the R^2^X value indicates that only a small portion of the receptor information (15%) is useful for predicting the therapeutic response.

The variable importance of projection (VIP) score was calculated to find which variables, across the entire PLSR model, most significantly contribute to determining the response to combination therapy (49). VEGFR1, CD36 and CD47 levels on tumor cells were found to be important for explaining the response, as they have VIP scores higher than one (***Fig. 5A***). The two nonlinear terms of tumor cell receptor numbers are also identified as important variables. We found that all of the receptors on tumor endothelial cells have VIP score lower than 0.5, indicating that tumor endothelial cell receptor expression was not as influential as receptor expression on tumor cells in predicting the response to combination therapy. Plotting the values of the receptors with VIP scores greater than one versus the response to combination therapy shows that lower VEGFR1 expression, lower CD47 expression and higher CD36 expression on tumor cells correlate with higher response (***Fig. 5B-D***). This indicates the patients with low tumor cell VEGFR1 or CD47 expression or high tumor cell CD36 expression are likely to show a stronger response to combination therapy than others. The associations between response and other receptors are not pronounced (***Supplementary Fig. S7***). The therapeutic response is also related to the two nonlinear terms in a monotonic fashion (***Fig. 5E-F***). Interestingly, although the response to treatment exhibits a trend that correlates with certain individual receptors, the data highly deviates from the LOWESS smoothing of the data (***Fig. 5, black lines***). This indicates that none of the 16 receptor variables can accurately predict the response alone, and the variability in the expression of a single receptor cannot account for all of the variability in the response to combination anti-angiogenic therapy. Hence, the ratio of receptor numbers is a valuable quantity in predicted the response to combination therapy.

**Figure 5.**
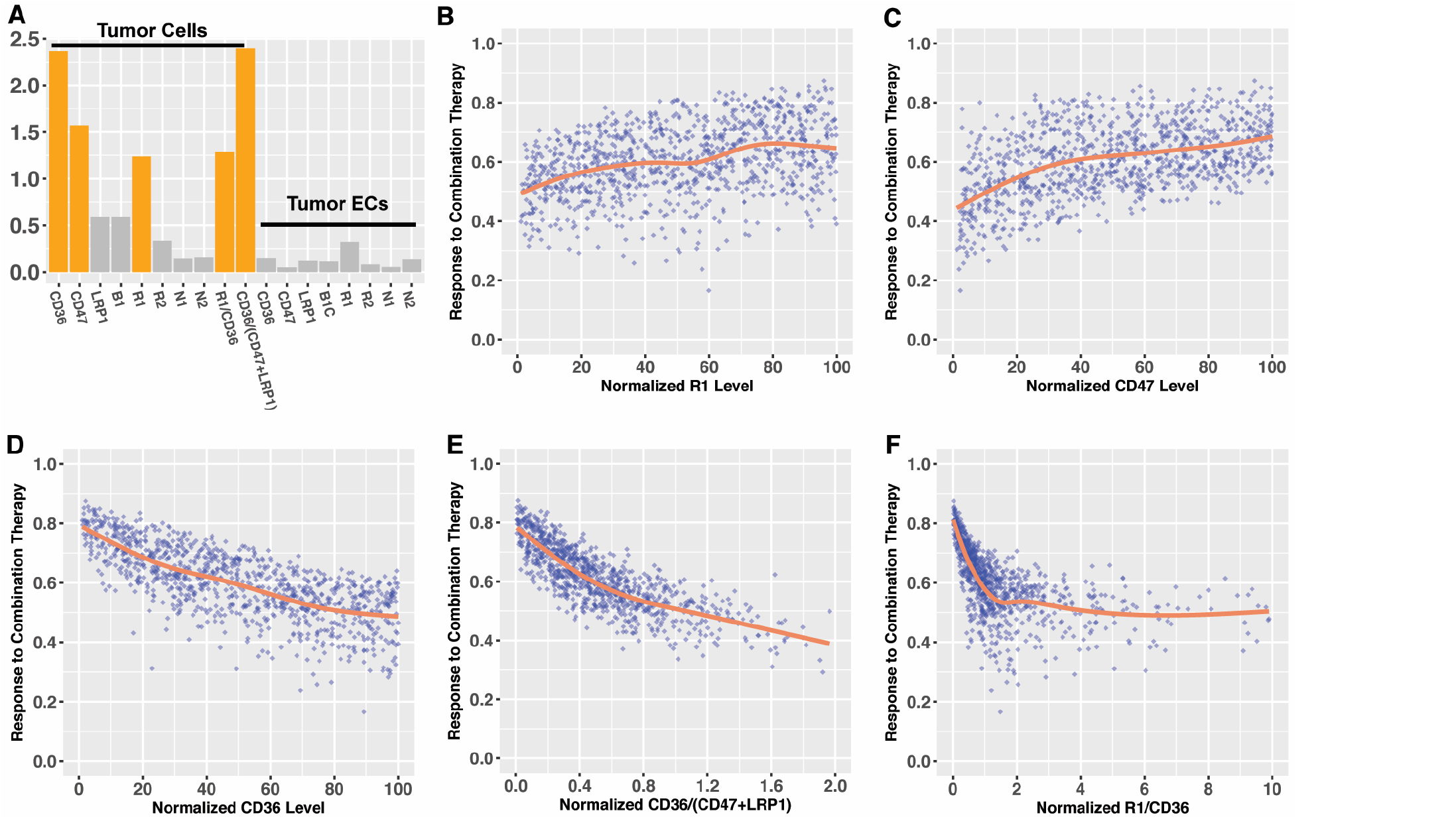
The association between tumor angiogenic receptors and the response to combination anti-angiogenic therapy. (A) The calculated variable importance of projection (VIP) score estimated by the PLSR model. VEGFR1, CD36 and CD47 on tumor cells have VIP scores higher than one (dark grey bars), indicating that they each significantly contribute to the response to treatment, the fold-change in the angiogenic ratio in the tumor. (B-F) The associations between normalized receptor level and the response to combination therapy. The orange line is the locally weighted scatterplot smoothing (LOWESS).

## Discussion

We present a systems biology model of the angiogenic balance in breast cancer patients. We significantly expand upon our previous work (22), extending the single compartment model to three compartments. This whole-body model allows us to account for the PK characteristics and distribution of anti-angiogenic drugs, which was not possible with our previous single-compartment model. This new model enables us to investigate the combination anti-angiogenic therapy targeting the VEGF and TSP1 axes. Additionally, the model parameters represent physiological and biological characteristics of cancer patients, allowing us to answer clinically relevant questions. Specifically, we investigated the impact of tumor receptor variability, one key aspect of inter-patient heterogeneity.

Overall, in the present study, we build our model based on biological knowledge and calibrate the model with both *in vivo* and *in vitro* data. We use the model to simulate anti-angiogenic therapy, generating new predictions and quantifying the impact of tumor receptor heterogeneity on the response to anti-angiogenic tumor therapy. Several model predictions are supported by published clinical observations, as discussed below.

This work uses a molecular-based PK/PD model to bridge the heterogeneous response of anti-angiogenic therapy with tumor receptor heterogeneity. Tumor angiogenic receptor heterogeneity has been observed in various settings. Qualitative information on the differential expression of angiogenic receptors from different tumor samples is reported in the Human Protein Atlas Database (26,27). Quantitative measurements also show the cell-to-cell variations in the density of VEGF receptors across different cell lines (24,25). This heterogeneity is thought to influence prognosis and response to treatment, which has been observed in various experimental settings. The VEGFR2 heterogeneity was proved to influence the response to anti-angiogenic cyclophosphamide treatment in a pre-clinical model (28). The inter-patient VEGFR1 and neuropilin heterogeneity has been associated with the response to bevacizumab treatment (29,30). Our modeling results provide a possible quantitative characterization of the mechanisms leading to the clinically observed heterogeneous responses to combination anti-angiogenic therapy (3,8,31), complementing the pre-clinical and clinical studies. The model predicts that after one cycle of treatment, the fold-change of the angiogenic ratio in tumor tissue varies widely depending on the tumor receptor levels. This indicates that tumor receptor heterogeneity could contribute to the heterogeneous response and should be considered when developing more effective and personalized anti-angiogenic therapy.

The characterization of the response in our study (the fold-change of angiogenic ratio) can be studied and measured in the experimental setting. The binding of angiogenic factors to their receptors initiates a cascade of intracellular reactions, including activation or phosphorylation of the receptors. The phosphorylated receptors can be measured (via flow cytometry or immunohistochemistry analyses of a tumor sample) and used to calculate the ratio of the pro and anti-angiogenic signals. Thus, the fold-change of angiogenic ratio reflects how the anti-angiogenic therapy changes the amount of phosphorylated pro-angiogenic receptors relative to the activated anti-angiogenic receptors, which together mediate tumor angiogenesis. Multiple experimental studies qualitatively explore the activated (phosphorylated) levels of the angiogenic receptors in the context of the TSP1-VEGF axis (45,51–54); however, quantitative data needed to validate our model are still missing. Therefore, a direct comparison with the clinical measurements is not possible. However, our modeling results are meaningful, a they identify important potential factors to measure (the tumor angiogenic receptors level before treatment), and can help design pre-clinical and clinical studies.

Our work helps identify biomarkers that predict response to anti-angiogenic treatment. The heterogeneous response to anti-angiogenic therapy is a major drawback for optimizing the combination therapy of bevacizumab and ABT-510, and generally, when considering all anti-angiogenic agents (3,31). However, despite substantial research (55), there is currently no reliable biomarker for identifying patients for which anti-angiogenic therapy will be effective. The PLSR analysis in our study showed that the levels of CD36, CD47, and VEGFR1 on tumor cells are potential biomarkers to select the patients who will likely respond to combination treatment. These receptors are shown to have a pronounced impact on the response: low expression of VEGFR1 and CD47 and high expression of CD36 all associate with high response to combination treatment. Interestingly, the relationship between VEGFR1 and the effect of bevacizumab has also been observed in clinical trial. Fountzilas *et al*. report that intra-tumoral VEGFR1 overexpression was an indicator for poor survival in metastatic breast cancer patients receiving bevacizumab and chemotherapy (56), which supports our finding and provides a qualitative foundation for our study. In addition to identifying these biomarkers, our study emphasizes the importance of the integration of information from multiple receptors to accurately predict the response. We showed that the expression of a single receptor is not able to fully account for the heterogeneous response to therapy. Instead, a comprehensive analysis that incorporates multiple biomarkers is more appropriate. In the experimental and clinical practice of biomarker discovery, it is commonly accepted that a single biomarker is not able to predict anti-angiogenic treatment efficacy (55). Our findings support this stance and demonstrate that a computational approach could help identify biomarkers by providing systematic understanding of the response to treatment. Specifically, the PLSR analysis indicated that the CD36/(CD47 + LRP1) ratio has the highest VIP score and is most important in predicting the fold-change in the angiogenic ratio in the tumor, indicating the importance of multiple receptors as a potential tissue biomarker.

Given the molecular detail of the model, we gain mechanistic insight into the model predictions. For example, we can understand how the individual tumor receptor populations change in response to anti-angiogenic therapy, insight that explains why VEGFR1, CD36, and CD47 are potential biomarkers. VEGFR1 has been supported as a possible *in situ* predictive biomarker for anti-VEGF agents in both preclinical and clinical settings (55–57). Our model predicts that the pro-angiogenic complexes involving VEGFR1 play an important role in the response to bevacizumab treatment: complexes involving VEGFR1 significantly decrease after the administration of bevacizumab while the levels of other pro-angiogenic complexes are relatively stable before and after bevacizumab treatment. The high expression of VEGFR1 can efficiently compete for VEGF, thereby limiting the benefits of VEGF neutralization by bevacizumab. On the other side, high expression of CD36 provides more available receptors for ABT-510 to bind to, which enables the formation of more anti-angiogenic complexes to shift the angiogenic balance. Since CD47 has a high affinity for TSP1, increasing the expression of CD47 will significantly elevate the baseline level of anti-angiogenic complexes. This means that the tumor has a higher baseline anti-angiogenic signal and consequently shows an attenuated response to the formation of more anti-angiogenic complexes by ABT-510. The insights generated by our model can guide future experimental or clinical studies of predictive biomarkers for anti-angiogenic treatment.

We do not include clinical response in this study due to lack of data needed to connect PK/PD data with clinical outcome. Thus, we currently cannot distinguish “complete response” and “partial response” in our model prediction at this point. However, we are able to get insights into outcomes from clinical trials involving anti-angiogenic agents from the predictions of our model at a qualitative level. ABT-510 failed in Phase II clinical trials for advanced renal cell carcinoma and metastatic melanoma (13,14). According to our model simulations, low CD36 expression on tumor cell could lead to the low observed response to ABT-510. Interestingly, the Human Protein Atlas shows CD36 is not detected in human renal cancer samples (in 10 out of 10 samples) and melanoma samples (in 11 out of 11 samples) (27). The heterogeneous expression of CD36 might be a possible reason for the failure of ABT-510. In the combination of bevacizumab and ABT-510 for advanced solid tumors, only 32% percent of the patients have prolonged stable disease for more than 6 months (17), which shows no evidence of being better than bevacizumab alone. Our model predicts that the difference between combination therapy and the best response of individual bevacizumab or ABT-510 treatment will become zero when the response to ABT-510 is very low. This means that the combination will not be superior to bevacizumab if ABT-510 alone is highly ineffective, which provides a possible explanation for the clinical observations.

Although we specifically simulate the combination of bevacizumab and ABT-510 in this work, our model can be adapted to study other anti-angiogenic agents that target VEGF and TSP1 signaling and can help explore different anti-angiogenic strategies. Given the many different anti-angiogenic agents (3,12) and possible drug regimens, our realistic model could be a valuable help in the search for an optimal combination anti-angiogenic strategy. Additionally, this model has practical interest, which can help circumvent difficulty in experimental studies. Traditional experiment-based study of comparing the effects of combination therapy to individual agents is limited by the difficulty of accumulating enough data to make meaningful comparisons. In contrast, our model can easily simulate multiple anti-angiogenic treatment strategies and quantitatively evaluate the added benefit of combination therapy as compared to single-agent therapy. In this study, we show that the combination of bevacizumab and ABT-510 leads to a stronger effect than administering either agent alone, in all of the sampled cases. However, the combination index revealed that the enhanced effect does not mean a strong synergistic effect. Our model would be a help to understand the synergism between two drugs and to design the combination therapy that achieves high efficacy with low drug dosage.

We acknowledge some limitations of the model that can be addressed in future studies. In this study, we use the angiogenic ratio to represent the angiogenic state, assuming that each angiogenic complex equally contributes to angiogenesis. However, it is known that the angiogenic receptors have unique functions and influence angiogenesis in different ways. For example, VEGFR1 primarily modulates blood vessel angiogenesis by ligand-trapping and receptor dimerization, while VEGFR2 is the predominant receptor that promotes pro-angiogenic VEGF signaling pathways (58,59). On the anti-angiogenic side, CD36 and CD47 are reported to inhibit angiogenesis by antagonizing survival pathways and activating apoptotic pathways (60). This knowledge of the relevant intracellular signaling pathways can be combined with our model to study specific functional responses to anti-angiogenic therapy. To date, several multiscale modeling studies have linked PK/PD models with the downstream signaling and cell-surface reactions (61,62). We can implement a similar approach, combining our model with downstream signaling models (63–65) to enable broader application of our model. Secondly, it is worth noting that there are alternate ways of characterizing the magnitude of the response to anti-angiogenic therapy. We focus on the angiogenic ratio, where the balance of pro- and anti-angiogenic factors has been experimentally shown to be a more accurate approach to study angiogenesis than analyzing the level of angiogenic factors individually (66). We further calculate the fold-change of the angiogenic ratio to characterize the response to anti-angiogenic treatment, while the area under the angiogenic ratio curve for the tumor compartment could be an alternative way to quantify the response to treatment. In that case, the response to ABT-510 is more substantial due to the strong shifting of the angiogenic ratio in the short time after each bolus (data not shown). However, the conclusions of our study remain unchanged. Another metric to consider is whether anti-angiogenic treatment shifts the angiogenic balance in tumor to the level observed in healthy tissue, related to vessel normalization (67). Since a universal definition of the response to anti-angiogenic therapy is still missing, the fold-change of the angiogenic balance of pro- and anti-angiogenic receptor complexes remains an important indicator, which this research is well suited to address. The primary focus for this study is the tumor receptor heterogeneity, and we do not investigate other possible inter-patient variations in this work, such as variation in the drugs’ the PK properties, different tumor sizes, or variable secretion of the angiogenic factors variation. Currently, these variations are poorly quantified and studied in pre-clinical and clinical experiments. Using our model as a tool to understand the impact of these other aspects of heterogeneity might be a great interest for future research.

### Concluding thoughts

Our model illustrates the effect of combination therapy of bevacizumab and ABT-510 on changing the balance between two opposing angiogenic signals. The model provides a quantitative description of the impact of tumor receptor heterogeneity on the response to combination anti-angiogenic therapy and aids in the discovery of predictive biomarkers. We expect that the insights generated by our model predictions will shed light on previously obscure clinical observations and that the model will be used to facilitate the optimization of new clinical trials.

## Acknowledgements

The authors thank members of the Finley research group for critical comments and suggestions.

## Supplementary Figures

**Figure S1. The comparison of predicted and measured pharmacodynamics of bevacizumab and ABT-510 in cancer patients**. The dots are measured data in clinical trials using bevacizumab (38) or ABT-510 (40) as single-agent treatment. The solid lines are predictions from the whole-body systems biology model.

**Figure S2. The sensitivity indices of model parameters**. We report the eFAST sensitivity indices for three groups of parameters: (A) receptor numbers; (B) inter-compartmental transport rate; and (C) the kinetic parameters.

**Figure S3. The effect of anti-angiogenic therapy on angiogenic complexes in tumor**. The left panel shows the changes of absolute concentrations of the angiogenic complexes. The right panel shows the percentages of individual angiogenic complexes relative to the total. (A) Bevacizumab promotes the decrement of pro-angiogenic complexes involving VEGFR1. (B) ABT-510 promotes the formation of anti-angiogenic complexes involving CD36. (C) Combination therapy decreases pro-angiogenic complexes and increases anti-angiogenic complexes.

**Figure S4. The dynamics of VEGF, TSP1, and the angiogenic ratio with combination therapy**. The solid line indicates the average of the simulation results of 1,000 sampled tumor receptor profiles. The shaded area shows the 90% confidence interval.

**Figure S5. The fold-changes of the angiogenic complex with combination therapy**. The numbers of the various receptor complexes change following combination treatment. Dark grey area indicates a fold-change higher than 1, and light grey denotes a fold-change lower than 1.

**Figure S6. The comparison of the predicted responses from the PLSR model and the mechanistic model**. Our optimal PLSR model has two principal components and shows good performance (Q^2^Y = 0.94; R^2^Y = 0.93). The dashed line represents a 100% match between predicted values from the PLSR model and the responses predicted by the three-compartment systems biology model.

**Figure S7. The association between angiogenic receptors and response to combination anti-angiogenic therapy**. The normalized receptor level is plotted versus the response to combination therapy (fold-change in the angiogenic ratio for tumor tissue) for all 16 receptors varied in our simulations. The orange line is the locally weighted scatterplot smoothing (LOWESS).

## Supplementary Files

**Supplementary File S1**. Detailed description of BioNetGen model formulation and parameters.

**Supplementary File S2**. This zipped file contains the computational model used to generate the results presented in this paper: “TSP1_VEGF_Model.m”, the MATLAB *m*-file containing the whole-body TSP1-VEGF model; “run_baselinedynamics.m”, “run_MCsimulations.m”, the MATLAB scripts to run the baseline model and Monte Carlo simulations, respectively; and “findParamValue_Rec.m” the MATLAB script to randomly generate the receptor number.

